# A melancholy machine: simulated synapse loss induces depression-like behaviors in deep reinforcement learning

**DOI:** 10.1101/2024.06.01.596905

**Authors:** Eric Chalmers, Santina Duarte, Xena Al-Hejji, Daniel Devoe, Aaron Gruber, Robert McDonald

**Affiliations:** Mount Royal University, Calgary, Alberta, Canada; Canadian Centre for Behavioural Neuroscience, University of Lethbridge, Lethbridge, Alberta, Canada

**Keywords:** Major Depressive Disorder, Reinforcement Learning, Neuroplasticity, Monoamine Hypothesis, Psychedelics, Reward Prediction Error

## Abstract

Deep Reinforcement Learning is a branch of artificial intelligence that uses artificial neural networks to model reward-based learning as it occurs in biological agents. Here we modify a Deep Reinforcement Learning approach by imposing a suppressive effect on the connections between neurons in the artificial network - simulating the effect of dendritic spine loss as observed in major depressive disorder (MDD). Surprisingly, this simulated spine loss is sufficient to induce a variety of MDD-like behaviors in the artificially intelligent agent, including anhedonia, increased temporal discounting, avoidance, and an altered exploration/exploitation balance. Furthermore, simulating alternative and longstanding reward-processing-centric conceptions of MDD (dysfunction of the dopamine system, altered reward discounting, context-dependent learning rates, increased exploration) does not produce the same range of MDD-like behaviors. These results support a conceptual model of MDD as a reduction of brain connectivity (and thus information-processing capacity) rather than an imbalance in monoamines - though the computational model suggests a possible explanation for the dysfunction of dopamine systems in MDD. Reversing the spine-loss effect in our computational MDD model can lead to rescue of rewarding behavior under some conditions. This supports the search for treatments that increase plasticity and synaptogenesis, and the model suggests some implications for their effective administration.

**Significance statement:** Simulating dendritic spine loss in a deep reinforcement learning agent causes the agent to exhibit a surprising range of depression-like behaviors. Simulating spine restoration allows rewarding behavior to be re-learned. This computational model sees Major Depressive Disorder as a reversible loss of brain capacity, providing some insights on pathology and treatment.

## Introduction

### Reinforcement learning: reward-based learning and behavior

Reinforcement Learning (RL) is a field of science that provides mathematical models of reward-based learning and decision-making. With its origins in the disparate psychological and computational fields of trial-and-error learning and optimal control ^1^, it is now a field where artificial intelligence (AI) and brain sciences overlap: recent RL-based learning algorithms can play *Go* and *Starcraft* at superhuman levels ^2,3^, while psychologists and neuroscientists use RL models to understand animal behavior ^1,4,5^.

Temporal difference learning is an important branch of RL which models trial-and-error learning. Temporal difference learning measures the difference between expected and actual rewards (the “temporal difference” or “reward prediction error”; RPE) each time the agent executes an action. The estimated value of that action is then adjusted in proportion to the RPE, according to a learning rate parameter. In the 1990s it was hypothesized that this model actually corresponds to learning mechanisms in the dopaminergic and striatal systems ^6,7^. This hypothesis has grown steadily, and today it is generally thought that the phasic activity of dopamine neurons in the midbrain signals reward prediction error (RPE) ^6^. It is believed that these phasic activity fluctuations influence plasticity in the striatum ^7^ (and possibly hippocampus ^8^ and prefrontal cortex ^9^) in a way that optimizes the organism’s expected or perceived value of particular actions, allowing it to learn rewarding behavior.

Deep Reinforcement Learning is a recent branch of RL which incorporates artificial neural networks: the networks are embedded within and control the actions of an artificial agent. In one possible implementation, the network accepts sensory input describing the environmental state, and the resulting activations of its output neurons represent the estimated values of various actions, given that state. Reward prediction errors experienced as the agent interacts with its environment are used to adjust the network’s internal connections in a learning process that improves the quality of the value estimates (thus maximizing the agent’s ability to make rewarding decisions), and generalizes knowledge between similar states (a critical ingredient in the ability to solve large problems and learn complex behaviors).

Because of the close analogy between Deep RL and biological reward-based learning, Deep RL is starting to be used in significant neuroscientific modeling and hypothesis creation, though much potential remains untapped ^10^. In particular, there is a tremendous but little-explored opportunity to use Deep RL to model *dysfunctions* of learning and behavior, including those associated with Major Depressive Disorder (MDD) ^11^. Artificial intelligence technologies have been used in diagnosis and treatment of depression ^12^, but a computational model of the disorder itself would be immensely valuable for MDD research: such models are cheaper and more ethical than animal models, can provide new ways of conceptualizing the disorder, and can produce new hypotheses for experimental researchers to investigate. This paper presents just such a model.

### Major Depressive Disorder - through a Reinforcement Learning lens

Major Depressive Disorder (MDD) is a complex psychological disorder. It is both highly disabling and highly prevalent: according to the National Institute of Mental Health, over 21 million Americans over the age of 18 have experienced a Major Depressive Episode over the last year ^13^. According to the DSM-5, symptoms of MDD include anhedonia, withdrawal from social activities, loss of interest, and increased indecisiveness ^14^. The ICD-10 categorizes recurrent depressive disorder as repeated episodes of depressive symptoms, including a decrease in overall mood and a reduction in the capacity for enjoyment of personal interests ^15^. Overall, MDD involves a general decrease in goal-directed behavior and productivity.

MDD is complex, heterogeneous, and not completely understood, but has been conceptualized in various ways. As it happens, some prominent conceptions involve or overlap with Reinforcement Learning and could be modeled using RL or straightforward extensions of RL. These conceptions of MDD include:

#### Monoamine deficiency

The “monoamine hypothesis” - the idea that MDD is primarily a dysregulation of monoaminergic systems (dopamine, serotonin, etc) - has been used for many years and inspired a variety of antidepressant drugs. Early generations of tricyclic antidepressants (TCAs) and monoamine oxidase inhibitors (MAOIs) eventually gave way to selective serotonin reuptake inhibitors (SSRIs), which can take several weeks to work but have fewer side effects ^16^. Amphetamines have sometimes been used ^17^ or paired with other drugs ^18^ because they increase dopamine levels. The downregulation of dopaminergic systems in MDD has been well documented ^19^ and is of particular interest from a Reinforcement Learning perspective: if dopamine signals reward prediction error, then disruption of that signal may also disrupt learning.

#### Increased temporal discounting

Temporal or delay discounting refers to the decrease in the perceived value of a future reward as the time it takes to reach the reward increases. Higher discounting rates mean a stronger preference for immediate reward over a delayed reward. It has been hypothesized that tonic activity of dopamine neurons affects discounting ^20,21^, and an increase in delay discounting has been associated with various psychiatric disorders, including bipolar disorder, bulimia nervosa, borderline personality disorder, and ADHD ^22,23^, and larger discounting rates are observed in depressed individuals relative to controls ^24,25^. This altered discounting may be a reflection of particular symptoms of MDD, such as feelings of hopelessness ^24^, or related to the impaired episodic future thinking that is observed in depressed individuals ^26^. While MDD is generally associated with increased discounting, it should be noted that the discounting phenomenon is complex and may vary with age and environmental factors ^27–30^.

#### Faster learning from negative experiences

Depressed individuals appear to have a heightened response to punishment or negative experiences and a blunted response to reward or positive experiences, both neurally and behaviourally ^31–33^. Such imbalances in feedback processing have been linked to altered dopamine signaling ^34^ and are also affected by SSRIs ^35^.

Anhedonic severity seems to correlate with attenuated processing of positive information in the striatum and enhanced performance in avoiding losses ^36^. However, reinforcement learning models with dual learning rates (separate learning rates for rewards and punishments) have not always detected faster learning from punishments when fit to behavioral data ^37–39^.

#### Altered exploration/exploitation balance

MDD is associated with altered exploration-exploitation tradeoff: the problem of deciding whether to exploit a trusted, rewarding option or explore other options that may potentially yield better outcomes. Depressed individuals exhibit an increase in stochastic choices and exploratory behavior ^40–42^, and this has been interpreted as the result of a decreased sensitivity to the value of different options ^11,43,44^.

However, as with the impaired processing of negative versus positive feedback, some studies that apply computational RL models have found that exploration levels do not account for the differences between MDD patients and controls ^45,46^.

### A new model: depression as reduced brain connectivity

Depression has been shown to affect dendritic spine density. Changes in spine density depend on brain region, sex, and other experimental conditions ^47^, but depression is generally associated with decreased spine density, especially in the hippocampus and prefrontal cortex ^47,48^, and lower synaptic density seems to correlate with depression severity ^49^. Dendritic spines are small protrusions from neuronal dendrites that can contact and form a synapse with a neighboring neuron. Fewer spines therefore suggests reduced overall brain connectivity. Some recent work suggests that ketamine relieves depression by promoting plasticity and spinogenesis ^50,51^ and that spines formed after Ketamine treatment are necessary for sustained antidepressant effects ^52^. Furthermore, it has been suggested that conventional antidepressants (that are based largely on the monoamine hypothesis) may work not by modulating monoamine concentrations *per se*, but because they promote slow synaptogenesis ^53–55^, or because they promote cognitive reconsolidation in a more positive way ^56^. Observations like these have affected a recent shift in thinking, away from MDD as a monoamine system dysfunction, toward seeing MDD as a dysfunction of neuroplasticity ^57^.

### Towards a computational model of depression

In this paper we set up a deep reinforcement learning algorithm to simulate dendritic spine loss. Specifically, we apply a “weight decay” effect to all connections between artificial neurons, causing each to degenerate over time at a rate proportional to their strength (i.e. the strongest communication channels are affected most - see Methods). This simulates the effect of spine loss by impairing the connections between neurons (illustrated in figure 1).

**Figure 1.**
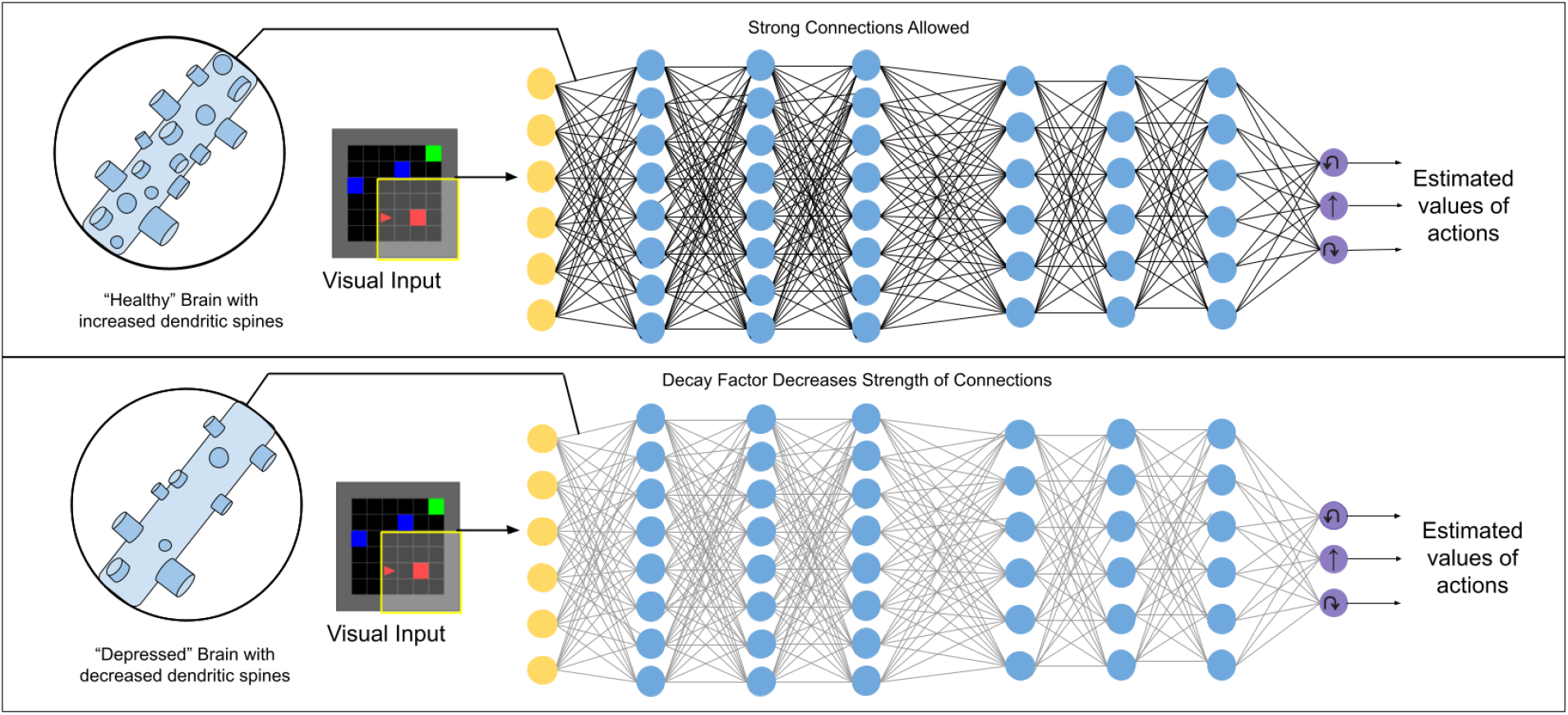
Illustration of the kind of deep reinforcement learning models used in this work. In an artificial neural network, the net communication channel between two neurons is represented by a single weight. We apply a weight decay factor to simulate the reduced connectivity effect of dendritic spine loss seen in depression.

We show, perhaps surprisingly, that this simulated spine loss is sufficient to induce a range of depression-like behaviors in the artificial agent, including anhedonia, temporal discounting, avoidance, and increased exploration. We further demonstrate that simulating alternate RL-centric conceptions of depression (reduced dopamine signaling, an artificially-increased discount rate, faster learning from negative experience, and altered exploration/exploitation balance) do not produce the same range of behaviors, showing that the simulated spine-loss model has face validity as a model of depression. We conclude by drawing several interesting hypotheses and implications from the model, including a new way to conceptualize depression, a speculative explanation for dopamine system dysregulation in MDD, and implications for treatments.

## Results

### A simulated “world”

Experiments in this paper use a simulated goal-seeking task illustrated in figure 2a. The simulation places an agent in a small room which it must learn to navigate. Three types of objects are present in the room: several “optional” goals, one “required” goal, and a “hazard”. Optional goals appear randomly throughout the room in each episode, and deliver a small reward to the agent when collected. The required goal always appears in the same location. When it is reached, the agent receives a large reward, the current episode ends, and the next begins. The hazard delivers a punishment (negative reward) to the agent when touched. At each step the agent chooses one of three actions: turn left, turn right, or move forward. The agent’s visual field only covers part of the room, so it must learn how to select appropriate actions to maximize reward, given the incomplete visual information. Both “healthy” and “depressed” agents (with simulated spine loss) were placed in this environment and allowed to learn behavioral strategies over a sufficient period of time.

**Figure 2.**
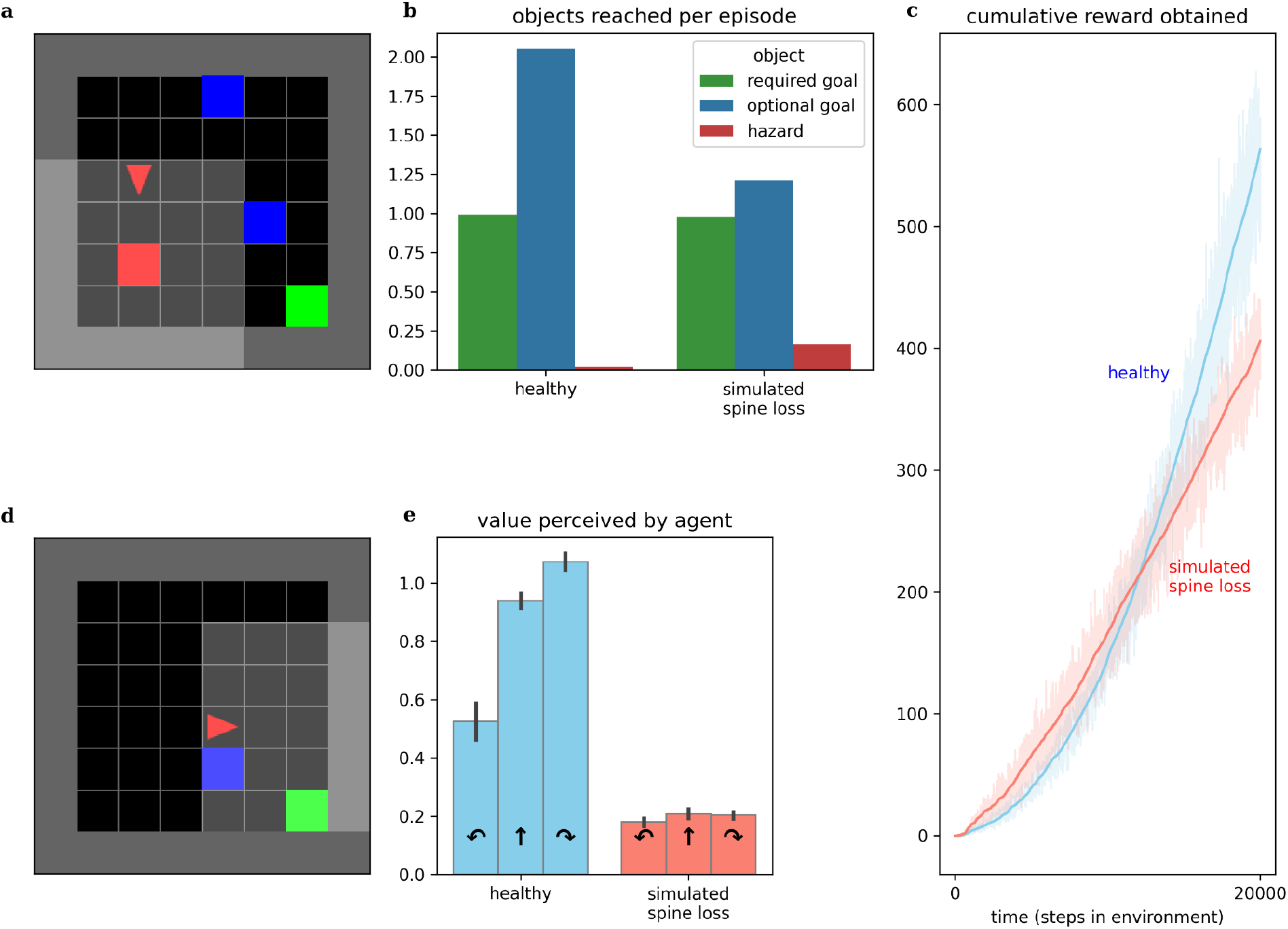
Comparing behaviors of simulated healthy and simulated spine loss agents. a) The simulated “world”. The agent (red triangle) must learn to navigate the room in search of the green goal. Blue boxes are optional bonus rewards that can be collected en route to the goal, and red boxes are hazards that bring a negative reward (punishment). b) The simulated spine loss agent still reaches the goal in each episode but collects fewer bonus rewards. c) The simulated spine loss agent learns a less-rewarding strategy than the healthy agent but learns it quicker, as seen by the time to arrive at asymptotic slope d) Contrived situation in which the agent may bypass the optional reward or take an extra step to collect it en route to the goal (reward optimal). e) agents’ perceived values for the contrived situation in d - the healthy agent’s perceived value of detouring through the optional reward is very high. The spine loss agent has much weaker preferences and a slight preference for bypassing the optional reward (an anhedonia-like effect).

While this simulation presents a goal-seeking problem involving spatial navigation, it should also be seen as a simple metaphor for the sequential decision-making of daily life. The optional goals represent opportunities like play, exploring interests, or socialization, while the required goal represents basic survival strategies like finding food or employment, which must be attended to every day.

### Simulated spine loss induces depression-like behaviors

#### Anhedonia

Anhedonia (loss of interest or pleasure) is recognized as a hallmark symptom of MDD by both the DSM-5 and the ICD-10 ^14,15^. Behavior of both healthy and spine-loss agents is illustrated in figure 2. Comparing the times required to reach asymptotic slope in the cumulative reward curves and comparing the asymptotic slopes themselves, we see that the “depressed” agents learn a simple strategy more quickly than the “healthy” agents, but the strategy learned by the “healthy” agents is more rewarding (see figure 2c). This is illustrated further by comparing the number of optional goals obtained per episode. “Healthy” agents collect almost two optional goals per episode, while “depressed” agents tend to collect only 1. If optional goals represent opportunities like play, exploring interests, or socialization, then these results mirror the loss of interest in these opportunities that accompanies depression.

Studies of MDD in animals often detect anhedonic (reduced pleasure-seeking) behavior using a sucrose preference test ^58,59^. Here, we instead place agents in the contrived scenario illustrated in figure 2d and observe their internal, perceived values. Assuming reasonably low reward discounting, the reward-optimal strategy would turn right to collect the optional goal en route to the required goal - requiring 1 additional action but obtaining both goals. The depressed agent chooses to bypass the optional goal - assigning a higher value to moving forward than to turning right. This is consistent with the results in figure 2b and suggests a simulated anhedonia. But the depressed agent also shows perceived values that are reduced categorically, revealing a general loss of interest underlying the anhedonic behavior.

#### Increased discounting

Altered discounting has been observed in MDD, both monetarily ^24^ and in response to basic rewards through simulations ^42^. Computational models provide the opportunity to directly expose the agent’s internal preferences and perceived values. We can infer an agent’s effective temporal discounting rate from these values. Figure 3b shows the simulated spine loss agent operating with a lower effective discount factor (more reward discounting). Surprisingly, this occurs *even though* all RL algorithms in these experiments used the same explicit discount factor setting. That is, the spine loss seems to induce an additional discounting effect.

**Figure 3.**
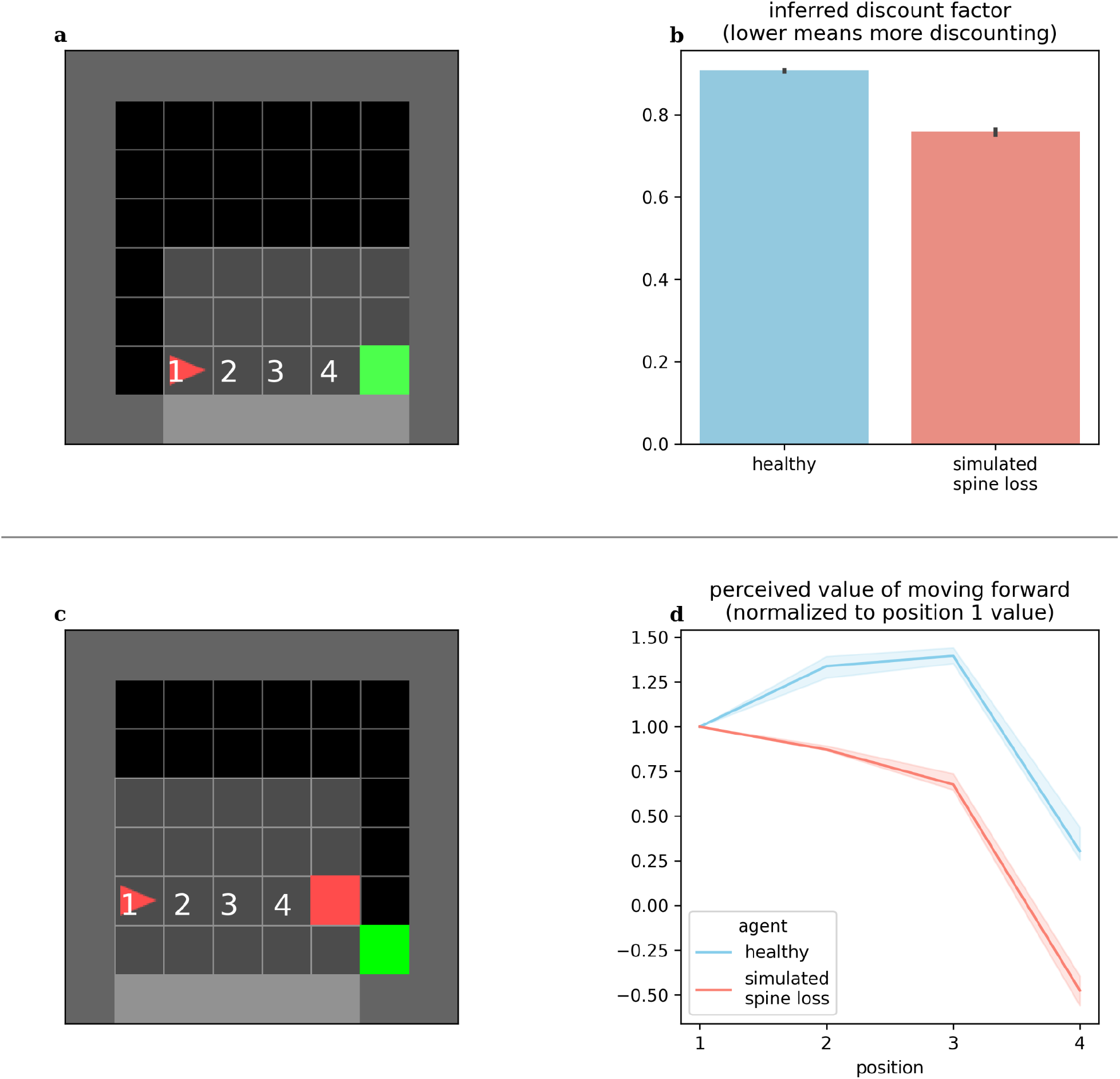
a) and b) agents’ effective discounting rates can be inferred by placing them progressively closer to the goal and measuring their perceived values. The spine loss agent operates with a lower effective discount factor (more discounting). c) and d) Contrived situation in which the agent is moved toward a hazard. The healthy agent’s perceived value of moving forward increases through positions 1-3 (moving forward from these positions brings the goal closer). Only in position 4 does the healthy agent’s perceived value of moving forward drop. For the spine loss agent this drop is generalized inappropriately to positions 2 and 3.

#### Avoidance

MDD involves maladaptive and increased avoidance behavior ^60,61^. We placed our computational agents in the contrived situation shown in figure 3c: the agent is slowly moved toward a hazard lying in front of the required goal. The healthy agent’s perceived value for forward motion increased through positions 1-3 (since this brings the agent closer to the goal). Only in position 4 does the value of moving forward drop - at that point the agent would prefer to go around the hazard. Conversely, in the depressed agent, the aversion to moving forward is (inappropriately) generalized to positions 2 and 3. Broekens et al. have suggested that such a decrease in perceived value within an RL model can be interpreted as analogous to fear ^62^.

### Simulated spine density modulates depression

By applying and then removing the dropout effect in the agent’s neural network, we can simulate the loss of dendritic spines and their subsequent restoration. Results in figure 4 show that depressive behaviors can be alternately induced and removed by applying or removing the connection weight decay factor. This seems to suggest that spine density, rather than being simply *correlated* with depression, modulates it. Interestingly, removing the simulated spine loss allows a return to the agent’s original behavior, but only after a brief drop in performance and a period of re-learning (this period is discussed further in the Discussion section).

**Figure 4.**
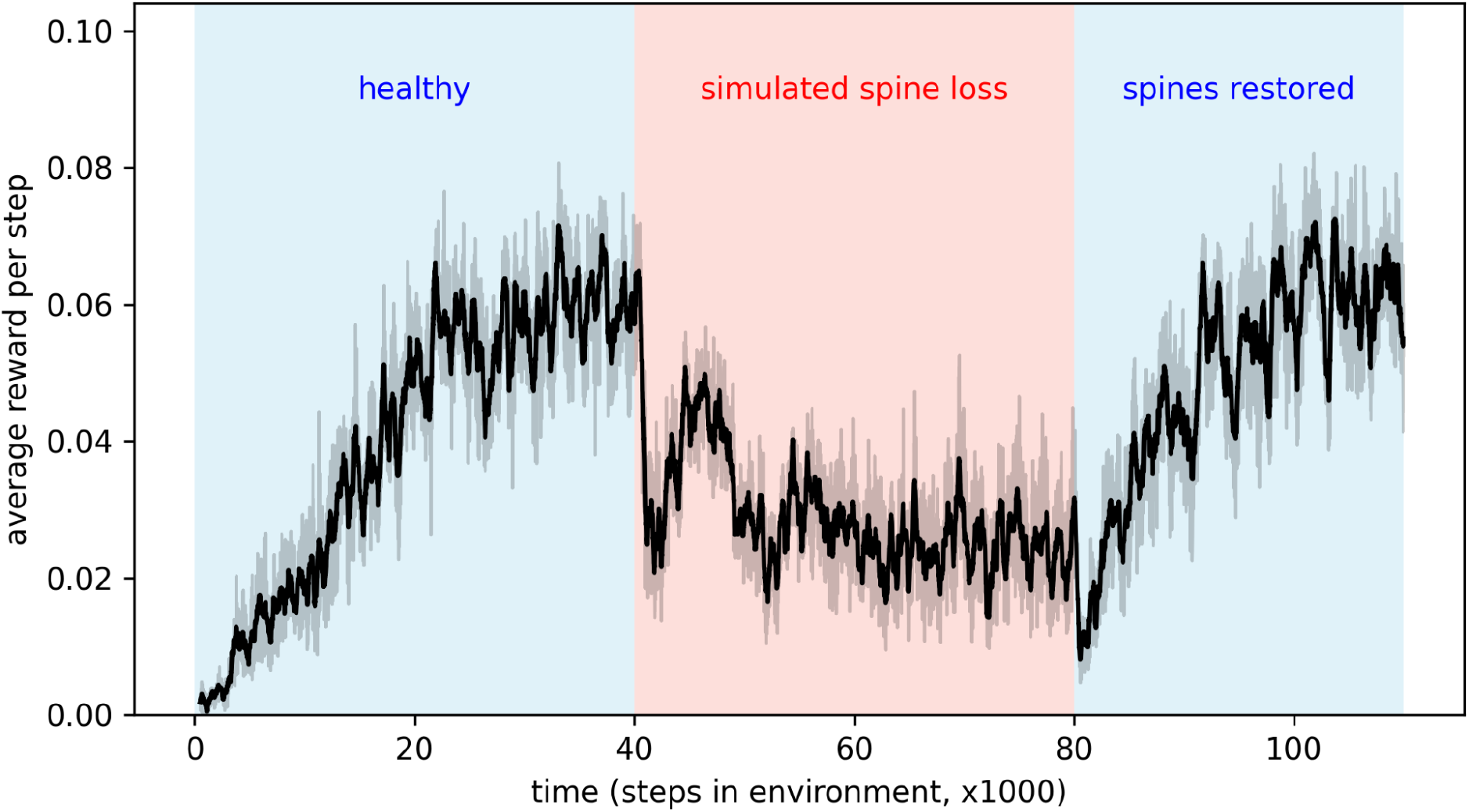
Applying simulated spine loss to a healthy agent causes it to revert to the simpler, low-reward depression-like behavior. Relieving the spine loss (restoring the spines) allows a return to the original behavior - after a short readjustment period. This may support the idea that spine density modulates depressed cognition and behavior.

### Simulated spine loss affects network plasticity and information content

Observing changes in the artificial neural networks over time, we see that a given reward prediction error generally induces less change in the network with simulated spine loss. Figure 5 shows a dramatic change in spine-loss-network weights early in learning (figure 2c suggests this represents the network quickly learning a basic survival strategy) but thereafter the spine-loss network shows less plasticity than the “healthy” network. Additionally, when the agents move toward the required goal, most neurons in the spine-loss network show activity which is highly correlated with this movement, suggesting that most of the network is dedicated to information about the required goal. In the healthy network, some neurons strongly correlate with the goal-directed movement, but others are not - suggesting those neurons are more concerned with other things (presumably information related to the optional goals and the hazard).

**Figure 5.**
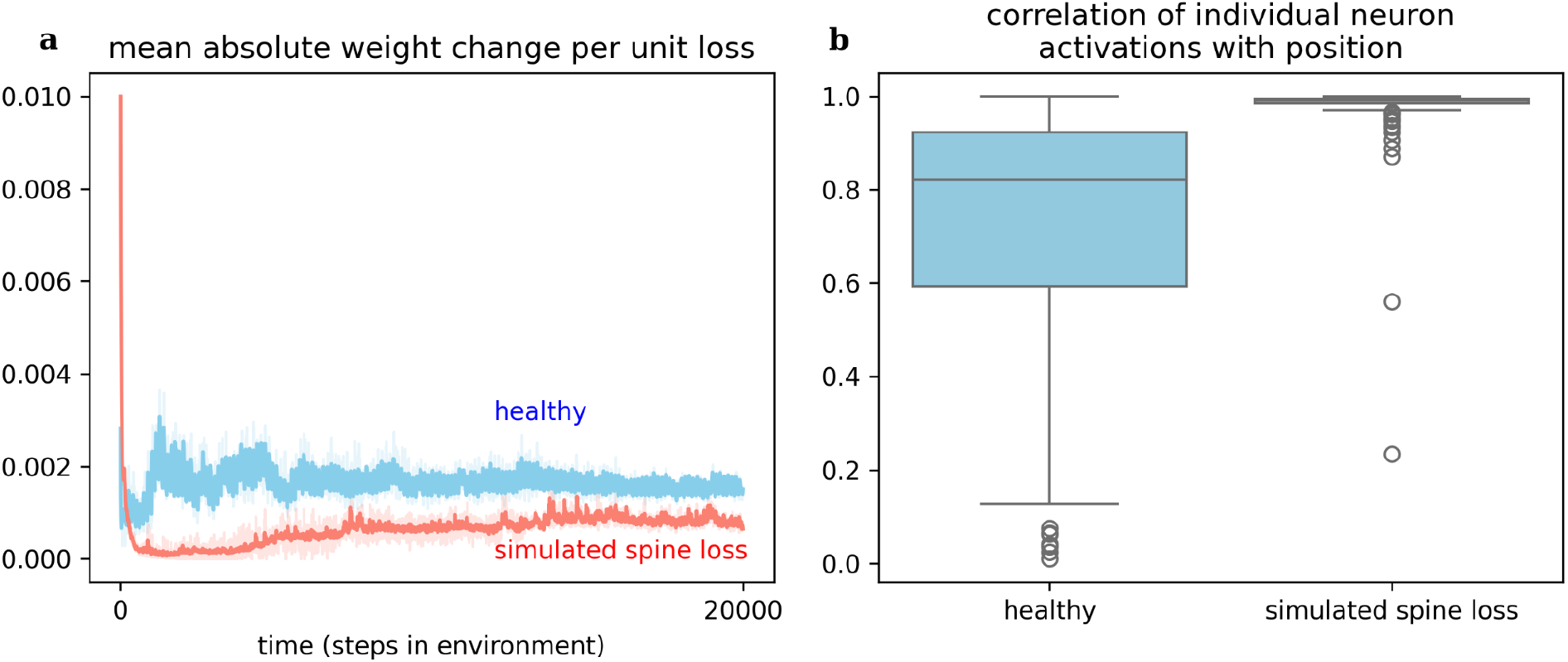
a) weight changes induced in the networks per unit loss. This measures simulated plasticity in response to the reward prediction error signal (i.e. delivered by dopamine). The spine loss network experiences dramatic alterations early in learning, allowing it to converge on a basic strategy quickly. The healthy network exhibits greater and sustained plasticity throughout learning. This effect may hint at an explanation for dopamine system dysfunction in depression. b) Most neurons in the network with simulated spine loss have activations highly correlated with proximity to the goal, indicating that almost the entire network has been used to store information related to the basic goal-seeking strategy. Neurons in the healthy agent’s network may store a greater variety of information.

### Simpler learning tasks are not impaired

The results above may give the impression that the spine-loss agent is *generally* impaired relative to the “healthy” agent. But figures 2c and 2b show the “depressed” agent converging on a strategy faster than the “healthy” agent - a strategy that does allow it to reach the goal every episode. To investigate this further, we create a simplified version of the environment by removing the optional rewards and hazards, leaving only the required goal. In this simpler task the spine-loss agent actually performs slightly better than the control agent (see figure 6). This seems to suggest that spine loss has a greater effect on more complex behaviors that require higher-order processing. One interpretation is that spine loss reduces overall network capacity, but leaves enough capacity for the network to learn simpler behaviors (see the Discussion section for further commentary).

**Figure 6.**
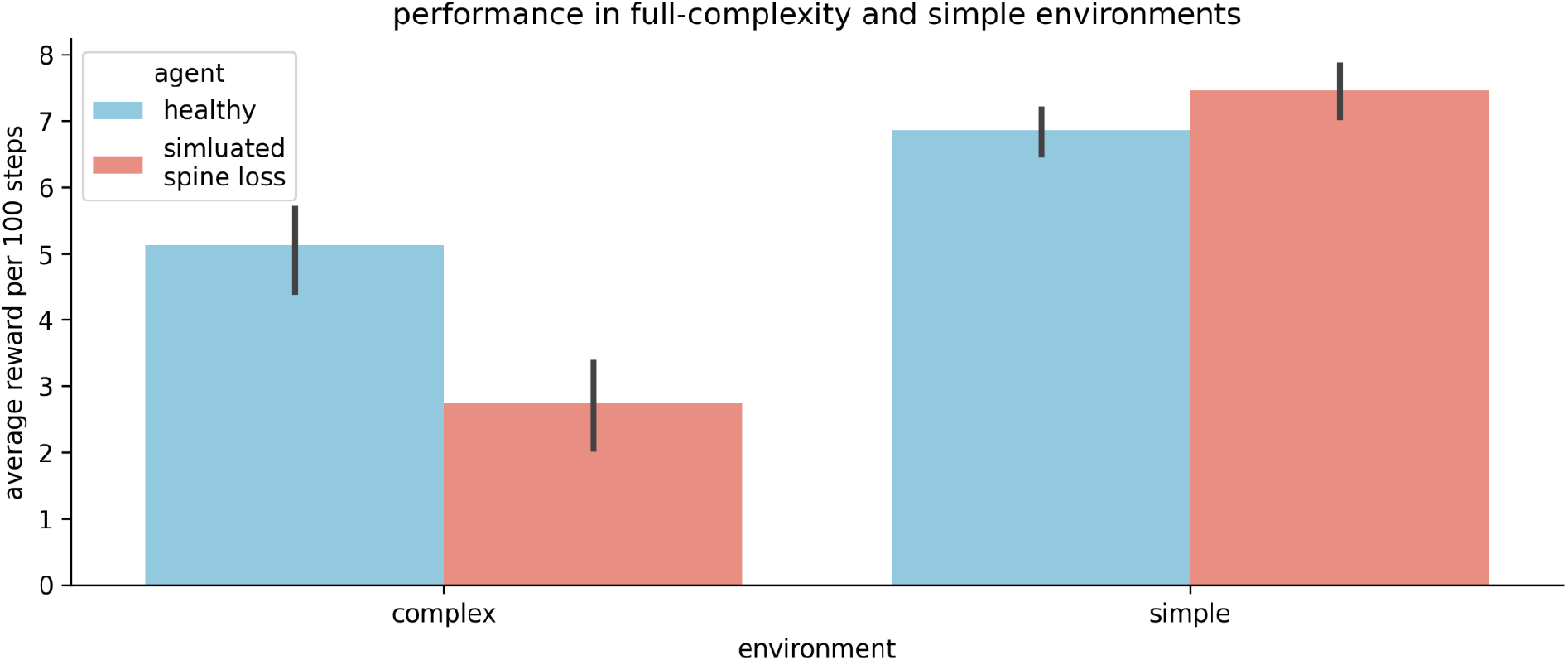
The “depressed” (simulated spine loss) agent is impaired relative to the “healthy” agent in the full-complexity task used throughout this paper. But in a simplified version of the task the impairment vanishes. Thus it seems that spine loss has a greater effect on complex behaviors that require higher orders of processing.

### A small amount of simulated spine loss is beneficial

The relationship between spine density and cognition / behavior is not well understood. And while depression is associated with decreased spine density, spines and synapses can be lost for many other reasons. During developmental pruning, for example, many synapses are eliminated from a young developing brain. Here we model adult brains in which the synaptic pruning process has already completed, but even in adulthood there is normal, ongoing turnover of a small number of spines and synapses ^63^.

Some turnover of connections is probably good, preventing stagnation and overfitting. This is well understood by machine learning practitioners, who often apply a small amount of weight decay to their artificial neural networks to improve learning and generalization ^64^. To illustrate this, we sweep through a range of weight decay settings from mild (representing normal, ongoing turnover of synapses) to extreme (representing pathological spine loss). As shown in figure 7, weight decay improves the agent’s performance up to a point: large amounts of decay cause depression-like impairments.

**Figure 7.**
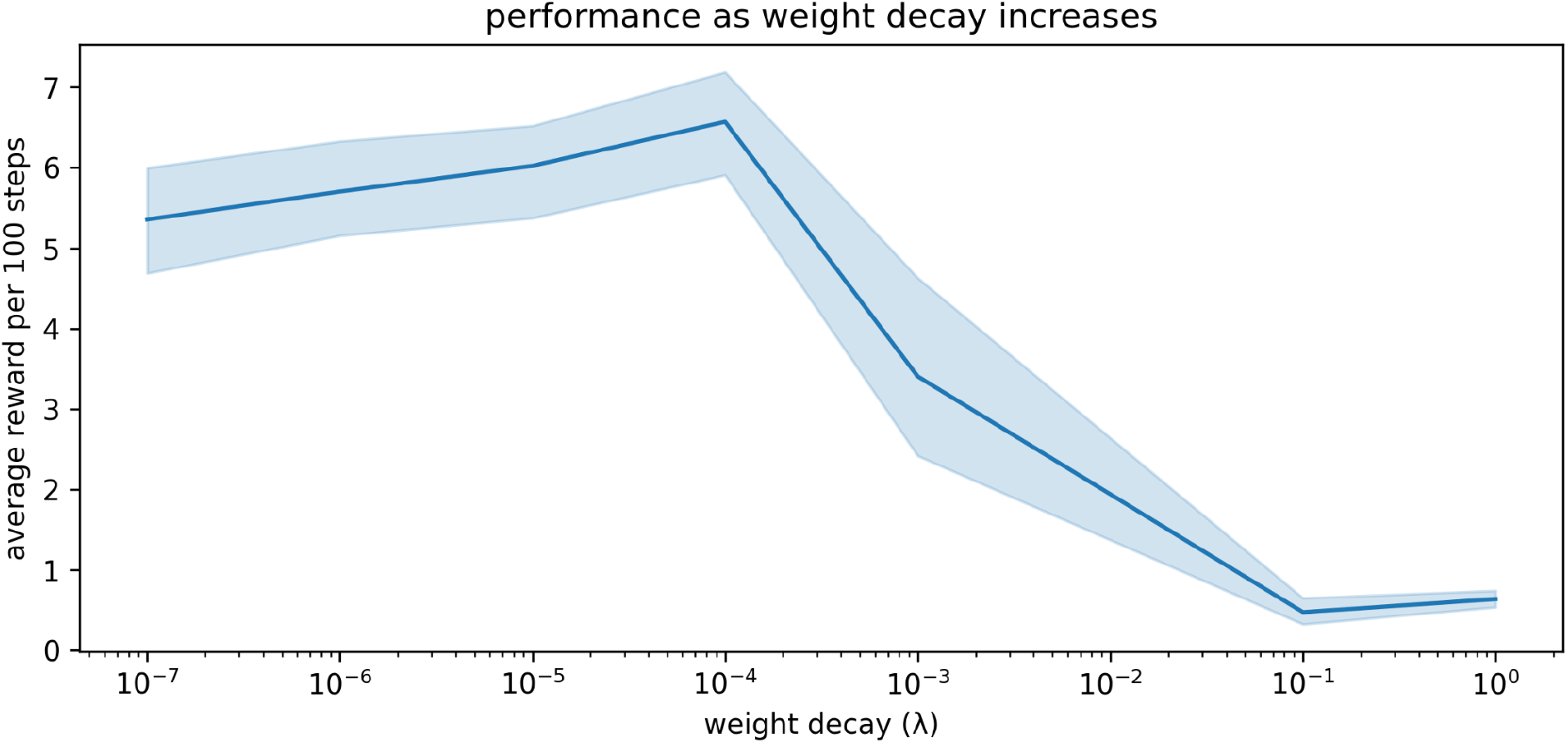
The relationship between weight decay and performance in our simulations. A small amount of weight decay (representing the normal, ongoing turnover of spines and synapses) is beneficial. More extreme weight decay causes impairment.

### Other conceptual models do not produce the same depression-like behaviors when simulated

We simulate the four other conceptions of MDD listed in the introduction: dopamine deficiency, increased temporal discounting, faster learning from negative experiences, and altered exploration/exploitation balance. Surprisingly, none produce all the same depression-like behaviors as simulated spine loss (table 1).

**Table 1.** The spine-loss model produces a wider range of depression-like behaviors than alternative conceptions of depression.

|  |  | Expected behavior |  |  |  |
| --- | --- | --- | --- | --- | --- |
|  |  | Anhedonia | Generalized avoidance/<br>fear | Increased discounting | More exploration |
| **Model of MDD** | Spine loss | ✓ | ✓ | ✓ | ✓ |
|  | Reduced reward-prediction-error signal |  |  |  |  |
|  | Faster learning from negative experiences |  |  |  |  |
|  | Higher discount factor |  |  | ✓ | ✓ |
|  | Increased exploration |  |  |  | ✓ |

## Discussion

### Implications for our conception of MDD

Computational models can help us fine-tune our conceptualization of a disorder. This computational model simulates spine density loss in a neural network, and surprisingly we find that this is sufficient to produce a variety of depression-like phenomena. These results encourage us to supplement our current understanding of MDD with the new (and complementary) view of MDD as a reversible loss of brain capacity.

If neural networks store information in connection weights or synapses ^65^, then, all other things being equal, a brain with more connections can store more information. Spine density loss reduces the number of connections available to the neural network’s learning process and thus reduces the capacity for storing information that supports choice strategies and behaviors.

Thus, the reversion or reduction to the simpler, basic survival strategy when spine loss is applied in figure 4: there is no longer enough room to store the richer, more rewarding, but more complex strategy. Similarly, if a depressed human brain only has enough room to hold the necessary behavior of going to work, then hobbies, socialization and other interests could be squeezed out of the neural network to some degree. This interpretation adds to (and seems roughly consistent with) the recent shift in thought away from MDD as a monoamine system dysfunction toward the idea of MDD as a dysfunction of neuroplasticity ^57^.

Future experimental studies could explore this idea by correlating neural activity with task performance in behaving animals. If the task is complex, such experiments should expect to see reduced task performance but clearer neural encoding of the main features of the task in depressed animals - a few results along these lines have been observed in other work ^66^, and would be consistent with our figure 5.

The idea of MDD as loss of brain capacity may also begin to reconcile some conflicting observations from computational literature. Implementing dual learning rates in reinforcement learning models has had mixed success in explaining depressed behaviors ^37–39^. Similarly, fitting exploration/exploitation parameters in computational models has had mixed success in accounting for depressed behavior ^38,45^. Our results may provide a clue for interpreting these findings. Our agents all used the same explicit temporal discount factor settings, yet the agent with simulated spine loss demonstrated additional discounting. That is, the reduced neural-network connectivity creates a discounting effect completely independent of the discount parameters used in conventional RL models - probably because the network must focus its limited capacity on learning how to seize nearby rewards. Similarly, faster learning from negative experiences and increased exploration may arise from reduced brain connectivity - in which case they would not be accurately accounted for by the explicit parameters used in conventional RL models. Future experimental studies could verify this conclusion by fitting our new deep-RL model to behavioral data.

### Dopamine system dysregulation as a cause of MDD? Or an effect?

Surprisingly, simulating spine loss in an artificial neural network is sufficient to produce a range of depression-like behaviors, but simulating reduced reward-prediction-error signaling by dopamine apparently is not. Still, dopamine system dysregulation is a well-documented feature of depression ^19^. How do we reconcile these facts?

The computational model suggests a speculative explanation for dopamine system dysfunction. Spine loss and reduced brain connectivity would limit the information-storage capacity of a neural network ^65,67^. Even basic survival strategies must fight for representation in a network with limited capacity, so there is less freedom for the network to adapt to the environment in arbitrary ways. Under these conditions, a dopamine spike unrelated to survival-critical events may go “unanswered” in terms of network changes. This explains the results in Figure 5, which show that the spine-loss-network’s response to reward prediction error (i.e. the dopamine signal ^6,7^) is muted relative to the healthy network. With fewer opportunities to act, the dopamine system may be thrown into dysregulation. Under this view, dopamine system dysregulation is not a *cause* of MDD; rather, spine loss and reduced brain connectivity cause *both* MDD and dopamine system dysregulation. It has been proposed that early damage to the hippocampus may lead to the altered forebrain dopamine operation seen in Schizophrenia ^68^ - perhaps the altered dopamine operation in MDD is, similarly, secondary.

There is a chance that thinking about MDD as a reversible loss of brain capacity could allow a wider (and very ambitious) reconciliation between the monoamine hypothesis and the newer neuroplasticity theory of MDD, by seeing neural network capacity as the link between the two. If the activation of 5-HT7 receptors promotes the formation of dendritic spines ^54^, and if those spines prevent depression-like symptoms by increasing network capacity ^53^, this may partially account for serotonin and tryptophan depletion causing depression-like symptoms in patients ^69^-the type of observation that originally led to the widespread use of SSRIs under the monoamine hypothesis ^57^. Clearly this is an inadequate summary of a complex (and contentious ^69,70^) topic. Still, limiting network capacity seems to account for a surprising range of depression-like behaviors - suggesting that various systems and interventions may exert their influence on depressive states *through* network capacity.

### Implications for MDD treatments

The model makes the hypothesis that spine density directly modulates depression. Many studies have associated spine density loss with depression, and some evidence for a causal link has been provided in the context of ketamine treatment ^52^. But in a recent review of psychedelic effects on neuroplasticity, Calder & Hasler conclude, “though changes in neuroplasticity and changes in cognition or behavior may occur simultaneously, whether neuroplasticity mediated those changes remains an open question for future studies to address.” ^71^. In light of this, the results in figure 4 are especially significant since they show the simulated spine-loss effect directly modulating depression-like behaviors. Though speculative, these results strengthen the argument for increased neuroplasticity and spine density as the sources of improved cognition and behavior, and support the search for treatments that promote spine growth.

This hypothesis that spine density modulates depression is consistent with the new thinking about MDD as a dysfunction of neuroplasticity, which has prompted research into treatments that upregulate plasticity. For example, the N-methyl-d-aspartate (NMDA) receptor antagonist, Ketamine, is theorized to promote synaptogenesis through one of several possible pathways that increase brain-derived neurotrophic factor (BDNF) levels and, ultimately, synaptic plasticity ^47,51,72–75^. Ketamine has indeed been observed to increase BDNF levels ^76^, increase spinogenesis and reverse dendritic atrophy ^52,75,77^, and produce fast and persistent antidepressant effects in patients and animal models ^51,78–81^. Psychedelics such as D-lysergic acid diethylamide (LSD), psilocybin, and dimethyltryptamine (DMT) have also been shown to promote neuroplasticity - possibly through a pathway that starts with serotonin 5-HT2A receptor agonism and involves stimulating BDNF production ^71,82,83^. They too can induce rapid antidepressant effects ^84–86^.

Furthermore, our model suggests some implications for the administration of such treatments. In figure 4, the period immediately after the simulated spine restoration is interesting because the agent does not immediately reacquire the high-reward behavior. Instead, the rate of reward actually drops, then slowly climbs back to pre-depression levels as the agent undergoes *general re-learning* of rewarding behavior for the environment. During this period the network is not simply augmenting the basic survival strategy stored during depression with new information about optional goals. Rather, the entire network is being reconfigured to make use of the new connections.

If a biological agent’s brain undergoes a similar (if much less dramatic) experience, then this period of time immediately after restoring lost spines is critical: experiences during this phase will influence what is re-learned. It is already known in the context of psychedelic treatments, for example, that the supportiveness of the environment during and after administration affects the treatment outcome ^71^. More research may be necessary to determine how to best use the “window of neuroplasticity” that remains open for a short time after treatment ^71^. We note that our artificial agent would not have reacquired the high-reward behavior if placed in an empty room after spine restoration: it requires the right environmental exposure to re-learn an appropriate strategy. This is similar to the conclusion drawn by Harmer et al. regarding conventional antidepressants: that they take effect slowly because they increase positive emotional processing, after which the patient must gradually re-learn their relationship to the world through accumulation of positively-processed experience ^56,87^.

More research is needed into the “window of neuroplasticity” effect generally, and based on these results, future experimental studies might examine the effects of environment and experience shortly after a plasticity-inducing treatment is administered.

### Limitations of the computational model, and opportunities for future work

As scientists, we often understand complex biological systems through *models* (e.g. animal models, mathematical / computational models, block diagrams, pictorial representations, etc). Every model is a sort of metaphor: it is *not* the complex target system, but it has something in common with - and therefore tells us something about - that system. Like all good models, our computational model abstracts away complexity in order to highlight some general principles. A real brain involves modularity, sparse and small-world connectivity, interactions between distinct structures, cortical columns, polysynaptic pathways, etc. Conventional artificial neural networks abstract away much of this complexity, and highlight the basic principle that rewarding behavior is learned through alteration of neuron-to-neuron connections. In reality, each of those neuron-to-neuron connections consists of a set of (sometimes redundant) synapses ^88^ that develop from among a large number of filopodia ^63^ across various functional zones of a dendrite ^89^. An artificial network abstracts away much of this complexity too, using a single weight to represent the *net* communication channel between two neurons. Such abstractions are allowing researchers to build tractable models of neural reinforcement learning; useful for testing high-level ideas and generating hypotheses ^10^.

Spine loss suggests fewer synapses and overall reduced connectivity between neurons. In an artificial neural network (where the usual approach is to represent the net connection between two neurons using a single weight) applying weight decay to each connection is a reasonable, abstract model of this reduced connectedness. The mechanisms underlying biological spine loss are a subject of debate ^90,91^, making it difficult to comment on the biological relevance of weight decay. It does seem that any mechanism which results in random pruning of spines or decreased probability of spine formation would, statistically, affect the likelihood of strong connections (involving many synapses) more than weak connections - an effect similar to that of weight decay, which degrades the strongest connections fastest. Similarly, mitochondrial dysfunction or otherwise altered energy dynamics within a neuron would seem to affect the strongest synapses with the highest metabolic cost. But until more is known, weight decay remains attractive mostly because it is reasonable and convenient.

We have shown the weight decay model already produces some interesting results. But even more insight may be available if future work can devise a more nuanced and biologically accurate way to simulate spine loss. For example, we have applied weight decay throughout the entire network, but in reality it seems spine density changes depend on brain region ^63,92^ and types of play experience ^93,94^. These kinds of effects are not included in our model, and so it can offer no insight into them. Making the simulations more biologically accurate in these ways may require significant re-thinking of the artificial neural network approach, but may create a new range of hypotheses. In the meantime, it may be useful to find other ways to compare the altered signaling in the artificial neural network with weight decay to experimental data.

All models exist on a scale from high abstraction to high detail: abstracting away complexity illustrates general “big picture” principles, which is appropriate for a first-of-its-kind study such as this one. We hope future work (our own and others’) will build more detailed models that complement (or indeed replace) this one.

## Methods

### Computational model

Our deep reinforcement learning agent is based on temporal difference (TD) learning, which supposes that an agent inhabits some state *s*, and can select an action to perform from some set of actions *A*. If executing that action leads the agent to a new state *s’*, and triggers some reward *r*, then the value of that particular experience can be formulated as:

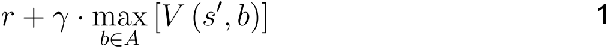

where 𝕪 is a discount factor (between 0 and 1) that discounts the value of future rewards relative to immediate ones. Note that *r* can be zero (no reward) or negative (a punishment). The value *in general* of executing action *a* from state *s* is then the average or expected value of all these individual experiences:

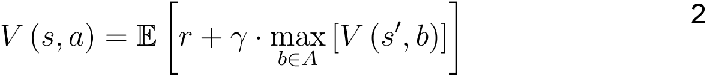

The learning process consists of updating the value estimates *V* after each experience in the world. A difference δ is computed between the experienced value and the current value estimate, and then used to adjust the estimate according to a learning rate α:

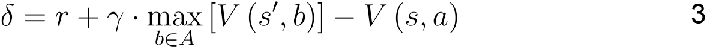

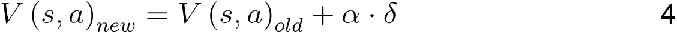

The difference δ is the reward-prediction-error that is thought to be signaled by dopamine neurons ^6,7^. For more information on TD learning see Sutton and Barto ^1^. Here we take a Deep Reinforcement Learning approach similar to that of Mnih et al. ^95^, in which an artificial neural network creates the value estimates *V*(*s*,*a*). This network accepts an input of 100 binary values (corresponding to the agent’s 5 x 5 visual field, times 4 object types that can exist at each location within the field). The network has a hidden layer of 10 neurons with tanh activation functions, and an output layer of 3 linear neurons whose outputs represent the values of turning left and right and moving forward.

The learning process now consists of tuning network connection weights such that the value estimates become increasingly accurate. For each experience, δ is computed per equation 3, the gradient of δ^2^ is calculated with respect to each network weight *wi*, and the weight is adjusted in the direction that will minimize δ:

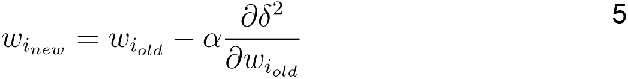

Note the analogy between equations 4 and 5. The weight decay effect is created by adding a sum of square weights, ½λ**w**2, to δ^2^ before computing the gradient. Equation 5 then becomes:

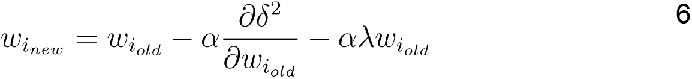

where λ controls how quickly each weight decays toward zero. A small amount of weight decay is often used with artificial neural networks to improve their performance and generalization ^64^, but here we use a more extreme weight decay setting to simulate significant spine loss.

See Supplementary Information for all parameter settings.

### Alternative model implementations

Our “healthy” agent was identical to the agent with simulated spine loss, except that the healthy agent used a weight decay setting of λ=0 (no weight decay). The variety of alternative computational MDD models shown in table 1 were implemented as follows:

#### Reduced reward-prediction-error signal

For this model, the reward prediction error signal δ was multiplied by a scale factor of 0.1 to simulate reduced dopamine signaling. Note that this scaling in the TD learning model is mathematically identical to using a smaller learning rate α.

#### Faster learning from negative experiences

This model multiplied all negative rewards by 2 and divided all positive rewards by 2 before doing network weight updates. Thus, negative rewards had a greater effect on those updates.

#### Higher discount factor

This model used a discount factor of γ = 0.5, rather than the γ = 0.9 used by other models. Per equation 1, this increases the discounting of future rewards.

#### Increased exploration

studies that use reinforcement learning to model animal behavior have often assumed a softmax action selection policy ^96–98^. Under this policy, the agent is assumed to select actions stochastically; actions with higher perceived value have a higher probability of selection:

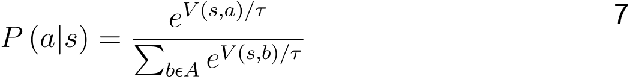

Here τ is a “temperature” parameter that sets the strength of the agent’s preference for exploiting the action with highest value, over exploring the other actions (for example, for high “temperature” values, the agent’s choice of action will be closer to random). Our “increased exploration” model uses this softmax action selection with τ = 1. This produces many more exploratory actions than the other models, which use an “ε-greedy” strategy of selecting the highest-value action with probability 1-ε, and a random (exploratory) action otherwise.

### Comparison with alternative depression models

Comparisons between the spine loss model and the other depression models are illustrated in figure 8, and were used to construct table 1. Models were considered to demonstrate anhedonia if there was no significant difference between the perceived values for turning right and moving forward in the experiment from figure 2d. Models were considered to exhibit generalized avoidance/fear if the perceived value of moving toward the hazard decreased prematurely, as in figure 3d. The spine loss model was the only model to exhibit anhedonia and generalized avoidance by these criteria, although the model with high discounting does exhibit lower perceived values in general (due to the extra discounting during evaluation of equation 3). Inferred discount factors were measured as in figure 3a/b. To evaluate relative differences in exploration rates, probability distributions were generated over actions (using equation 7). The distribution for the healthy agent was compared to those for other models using Kullback-Leibler divergence, for a number of randomly-generated world states. This approach uses the healthy agent’s probability distribution as a baseline, and measures deviation from that distribution in the other agents.

**Figure 8.**
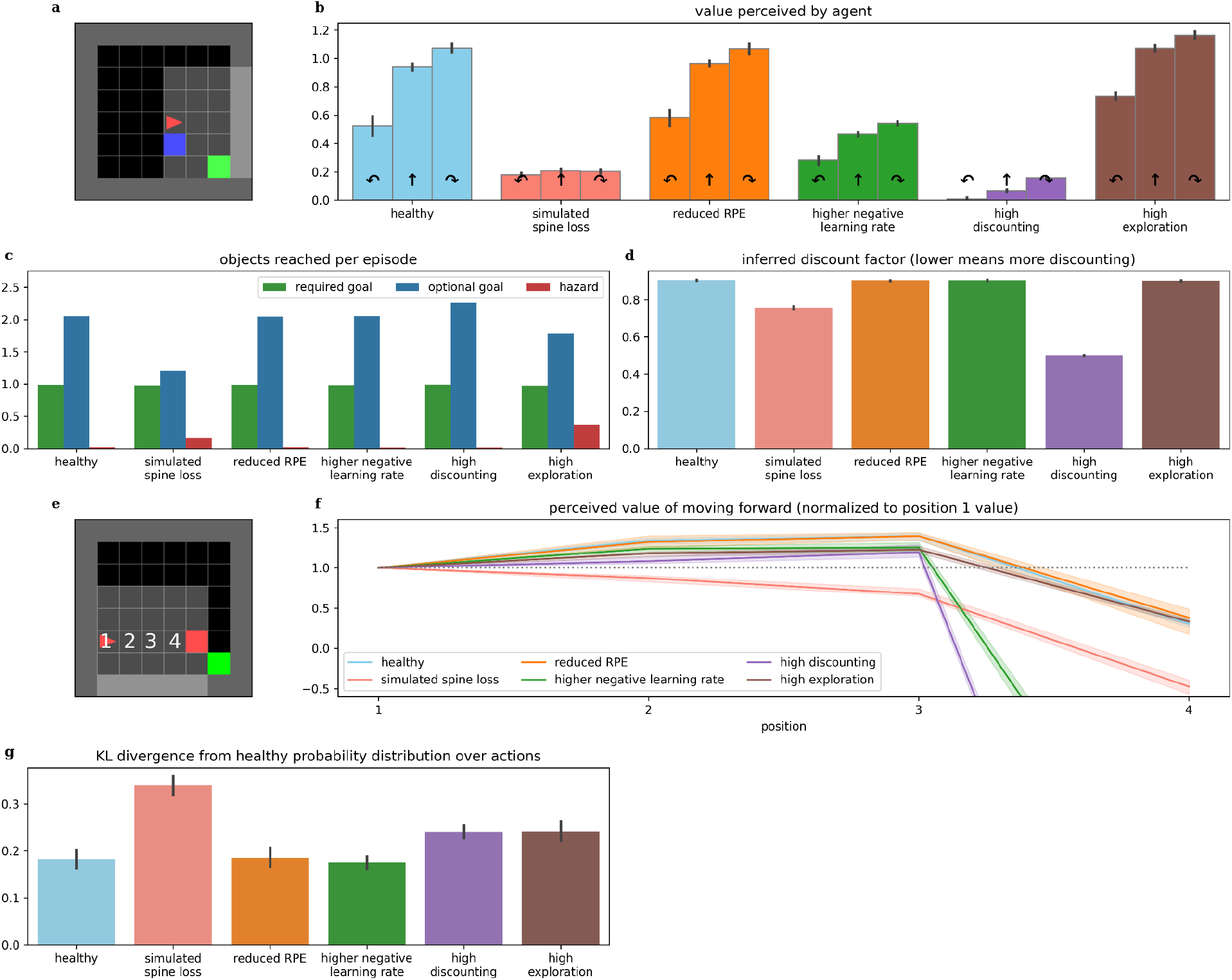
Comparing behaviors across alternative models of depression. b) The simulated spine loss agent is the only one that exhibits the strong anhedonia-like loss of preference in the contrived situation in a), although faster learning from negative experiences and increased discounting both cause a reduction in overall perceived values. c) The spine loss agent shows the most dramatic reduction in optional-reward-seeking behavior. The high-exploration agent shows a smaller drop in optional rewards due to the increased randomness in its actions. d) The agent with a high discounting parameter setting obviously exhibits higher discounting than other models. But the spine loss agent also shows increased effective discounting - despite having the same discount factor parameter setting as the healthy agent. f) The simulated spine loss agent is the only one exhibiting generalized-fear-like effects in the contrived situation in e). g) Kullback-Leibler divergence between probability distributions over actions, assuming a softmax action selection policy. All bars show the divergence from the probability distribution of healthy agents (“healthy” shows divergence between different healthy agents).

## Data availability

All code used in this work is available at https://github.com/echalmers/blue_ai.

## Acknowledgements

The authors gratefully acknowledge funding from Mount Royal University, and thank Jesse Viehweger and Ezzidean Azzabi for helpful contributions to this work.

## Author contributions

All authors contributed to experiment design and interpretation of results. EC, SD and XA performed literature review. EC executed experiments and collected results. EC, SD and XA prepared manuscript. All authors reviewed the manuscript.

## Additional information

The authors have no competing interests to declare.

## Notes

### Competing Interest Statement

The authors have declared no competing interest.

